# Highfold: accurately predicting cyclic peptide monomers and complexes with AlphaFold

**DOI:** 10.1101/2023.08.27.554979

**Authors:** Chenhao Zhang, Chengyun Zhang, Tianfeng Shang, Xinyi Wu, Hongliang Duan

## Abstract

In recent years, cyclic peptides have gained growing traction as a therapeutic modality owing to their diverse biological activities. Understanding the structures of these cyclic peptides and their complexes can provide valuable insights. However, experimental observation needs much time and money, and there still are many limitations to CADD methods. As for DL-based models, the scarcity of training data poses a formidable challenge in predicting cyclic peptides and their complexes. In this work, we present “High-fold,” an AlphaFold-based algorithm that addresses this issue. By incorporating pertinent information about head-to-tailed circular and disulfide bridge structures, Highfold reaches the best performance in comparison to other various approaches. This model enables accurate prediction of cyclic peptides and their complexes, making a step towards resolving its structure-activity research.

## 1. INTRODUCTION

Cyclic peptides have considerable potential as drug candidates with advantages over traditional small molecules.^1-3^ With the representative cyclic structure, those peptides preserve properties in terms of high binding affinity, selectivity, and specificity. Those peptide structures and related complexes are versatile tools to study activity and play key roles in pharmacology. However, predicting their monomer and complex structures is a long-standing challenge.

Traditional experiments can deliver excellent observation accuracy, but it often demands extensive time and effort^4-6^. Although there have been recent advances in approaches of Computer-Aided Drug Design (CADD) methodologies^7-14^, the computational prediction of cyclic peptide monomer structure still is mildly unsatisfactory. Those methods typically utilize physics-based energy functions and Machine Learning (ML) algorithms such as Monte Carlo sampling methods, indicating that they require an immense amount of time to sample for finding the lowest energy structures. As a result, these approaches generally can’t balance the trade-off between model accuracy and resource efficiency in predicting cyclic peptide folds. What’s more, CADD methods usually don’t take macrocyclic peptides(length>30) into consideration. Take Rosetta as an example^7^, its simple_cycpep_predict module is suitable for predicting small cyclic peptide (∼5-20 residues) conformation and it needs to run genkic_closure_attempts multiple times (from 250 to 50,000 could be accepted, depending on the input peptide) to find a closed conformation. As for predicting cyclic peptide complexes, AutoDock CrankPep (ADCP)^10^ is a docking tool that should be mentioned. Similar to Rosetta, this approach only can deal with cyclic peptides of up to 20 residues in length. Additionally, this method heavily relies on experimental structural information of target proteins to dock cyclic peptide structure generated by its algorithm. Namely, this approach doesn’t solve the cyclic peptide complex folds problem in a meaningful sense when lacking the target protein experimental 3D conformation. Considering the challenges in predicting cyclic peptides and their complexes, there is a particular dearth of a computational approach that can deliver high and reasonable predictions without relying on experimental structural data of the target proteins. Currently, DL-based methods such as AlphaFold^15^ and RoseTTAFold^16, 17^, have demonstrated remarkable accuracy in protein prediction and design, far surpassing the performance of traditional CADD methods^18^. It may be possible to use such models to make accurate predictions involving cyclic peptides.

While those deep learning networks have been considered a revolutionary change for accurate structure prediction^19-22^, in their current form it is not straightforward to predict cyclic peptide and their complexes. Until 2023, the study of Bhardwaj et al.^23^ demonstrated a model called AfCycDesign that adapted to predict the structures of cyclic peptide monomers, proving the feasibility of AlphaFold in dealing with cyclic folds. However, we found previously in this study of predicted models disregard the potential presence of disulfide bridges within the cyclic structure. Disulfide bridges play an integral role in cyclic peptide scaffolds, where they stabilize specific peptide conformations and significantly improve resistance to protease degradation^24, 25^. The absence of additional disulfide bridge constraint may lead this model to give unreasonable cyclic peptide folds, particularly in cases where there exist multiple disulfide bonds.

In this work, we propose a novel algorithm termed HighFold that both takes the head-to-tailed circle and disulfide bridges into consideration of AlphaFold and AlphaFold_Multimer, meanwhile allowing for the prediction of the cyclic peptide with different lengths. The core protocol that has enabled HighFold to predict cyclic structure is the bespoke N×N cyclic peptide matrix. The N is the peptide length. This matrix can indicate the relative position of residues of one peptide, establishing the circularization of cyclic peptides and connections for disulfide bridges. Notably, HighFold enumerates all possible disulfide bridge pairs assembles and creates their corresponding matrixes to AlphaFold/AlphaFold_Multimer. Then models give the top-5 results according to confidence values. This enumeration method can make a more precise and reliable prediction of disulfide bridge structures, alleviating the need for independent judgment by models in such cases.

As is shown in Figure 1, HighFold contains two models: HighFold_Monomer which depends on AlphaFold, and HighFold_Multimer which depends on AlphaFold_Multimer. The former focuses on the cyclic peptide monomer structure and the latter contributes to predicting cyclic peptide complex structural folds. Compared to the High-Fold_Monomer, the HighFold_Multimer gets the additional target protein input. To discriminate the cyclic peptide ligand and target protein structure features, we divide the input matrix into two parts: one is the default submatrix for the target protein and one is our novel submatrix for the cyclic peptide ligand. In our experiment, we adopt the dataset of AfCycDesign as a benchmarking to illustrate the RMSD comparison between HighFold_Monomer and it in monomer structure prediction. As for the evaluation of models in cyclic peptide complexes prediction, we chose the dataset of ADCP^10^ and implement the F_nat_ as the main metric.

**Figure 1.**
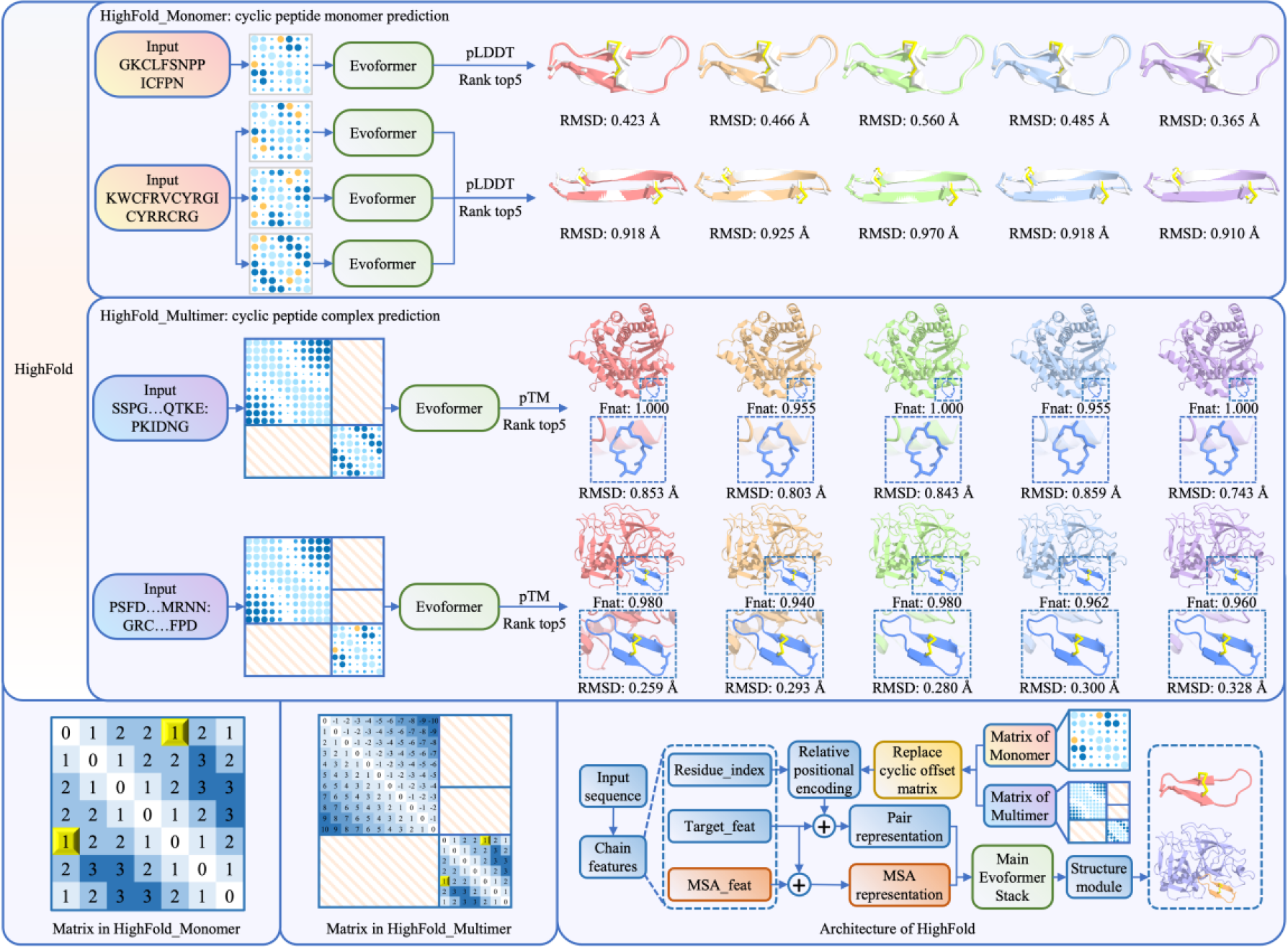
The workflow of HighFold. The HighFold encompasses a duet of modules: High-Fold_Monomer and HighFold_Multimer. In instances where the count of cysteine residues within the peptide input < 2, HighFold_Monomer only considers the head-to-tail structure and gives top-5 predicted cyclic peptide structure according to pLDDT scores. Conversely, when the number of cysteine residues ⩾ 2, HighFold_Monomer takes both the head-to-tail and disulfide bridge structures into consideration and enumerates all possible disulfide bridge pairs assembles, subsequently presenting top-5 predicted cyclic peptide structures in accordance with their pLDDT scores. The prediction process of HighFold_Multimer is similar to that of HighFold_Monomer. The differences between them are that the matrix of HighFold_Multimer has an additional protein target matrix and rank metrics is pTM rather than pLDDT score. The details about it can be found in the lower left corner and the total architecture of our model is displayed in the lower right corner.

To sum up, we propose an automated method, HighFold, to create predicted cyclic peptides and their complex folds by applying a novel input matrix. More importantly, to our best acknowledge, this is the first attempt that capable of predicting such cyclic peptides and their complexes’ structural folds involving head-to-tailed and disulfide bridge structures. As a DL-based tool for cyclic structures, this method shows great potential in accelerating the discovery of novel cyclic peptides and complexes.

## 2. RESULTS

### 2.1 The performance of HighFold_Monomer in cyclic peptide monomer structure prediction

In this work, we apply HighFold_Monomer and AfCycDesign which contain different matrixes to predict the structures of diverse cyclic peptides in the same experimental setting. From the data made available by AfCycDesign, we handpick 63 NMR structures with disulfide bridges. These structures span a diverse array of sizes, secondary structures, and sequences. Utilizing the capabilities of HighFold and AfCycDesign, we predict the three-dimensional structures for each sequence within the dataset. Subsequently, we assess the metric of backbone heavy atom RMSD to the corresponding experimentally determined structures. It should be noted that the pLDDT is adopted to rank predicted models in our work and we chose the top-1 predicted models for detailed analysis.

#### 2.1.1 RMSD comparison between HighFold_Monomer and AfCycDesign

Figure 2A shows the RMSD comparison between HighFold_Monomer and AfCycDesign in predicting cyclic peptide monomers. In general, the predictions emanating from the HighFold_Monomer framework exhibit remarkable proximity to the experimentally-determined structures with a median RMSD of 1.058 Å, and an average RMSD of 1.478 Å, respectively. In 37 out of 63 test cases, the predicted structures show backbone RMSD of less than 1.5 Å to the native structures. In only 5 cases, the predicted structure shows that the backbone RMSD of the natural structure is greater than 3 Å. Compared with the preceding method, HighFold improves the accuracy of prediction with an average of 0.259 Å, indicating the prowess of HighFold in cyclic peptide structure prediction. To further display the difference between those two models, we show the RMSD distribution in Figure 2B. The HighFold_Monomer has a deeper understanding of cyclic peptide structure and its RMSD results are centered largely in the bottom. What’s more, Figure 3 displays representative structures that HighFold_Monomer predicts correctly but AfCycDesign predicts wrongly. In predicting the 2mw0 (Figure 3A), AfCycDesign gains a good RMSD (0.990 Å). However, the predicted model of HighFold_Monomer can be closer to the native structure with the RMSD of 0.975 Å. In the task of prediting of 2m77, the gap between those two models is more evident and HighFold_Monomer is much better than AfCycDesign with the RMSD value of 1.530 Å.

**Figure 2.**
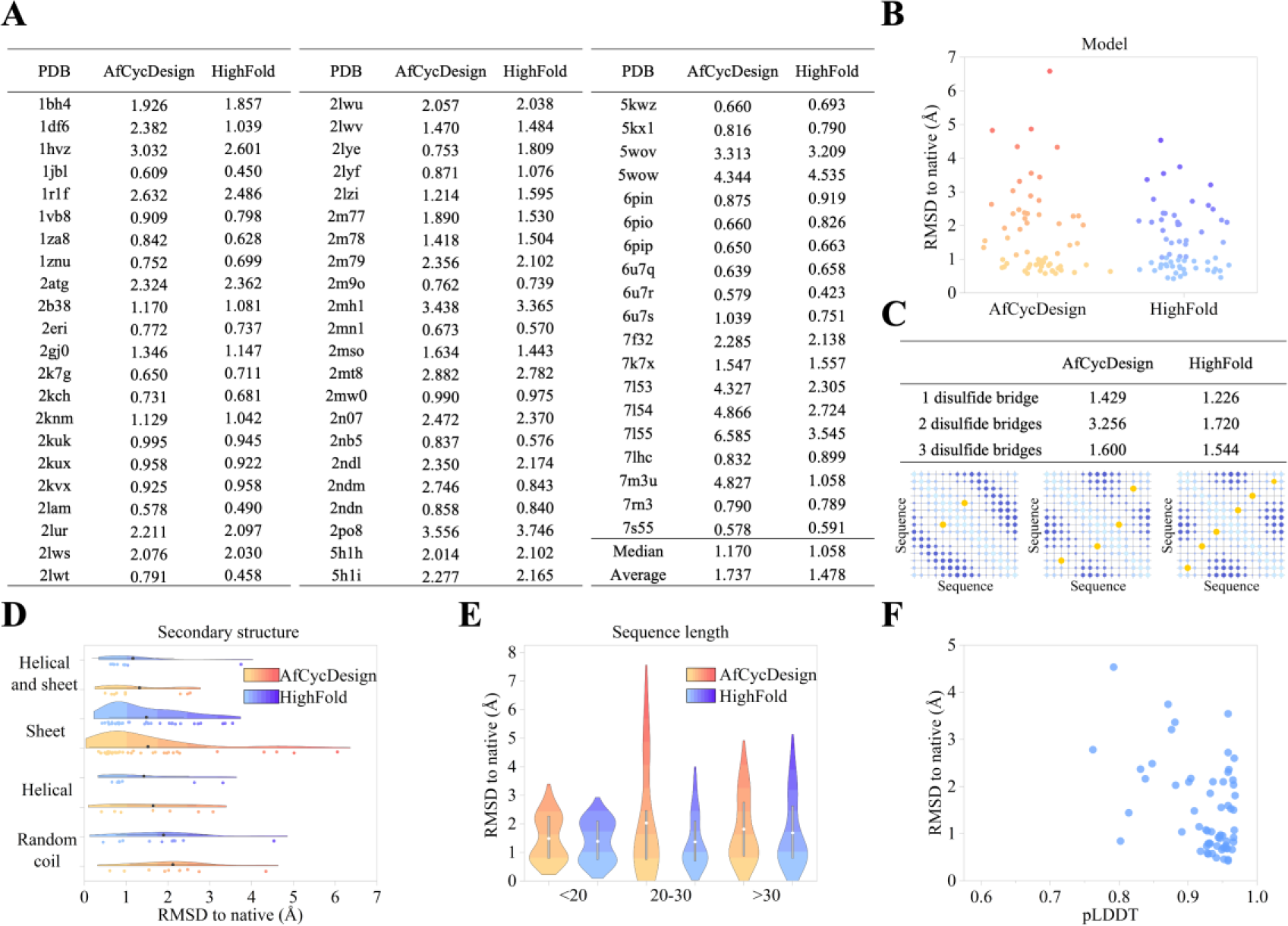
The results of models in cyclic peptide monomer structure prediction. (A) RMSD results of HighFold_Monomer and AfCycDesign. (B) RMSD distribution of HighFold_Monomer and AfCycDesign. (C) RMSD analysis of predicted disulfide bridge structures in HighFold_Monomer and AfCycDesign. (D) RMSD distribution of HighFold_ Monomer and AfCycDesign based on different secondary structure motifs. (E) RMSD distribution of HighFold_ Monomer and AfCycDesign based on input lengths. (F) The correlation between RMSD and pLDDT. Note that all data analyses are based on top-1 results.

**Figure 3.**
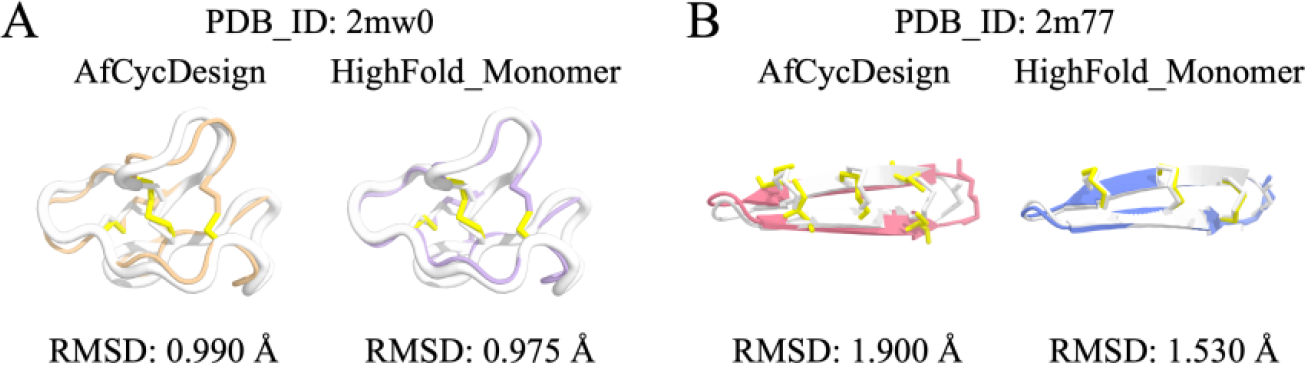
Structures that HighFold_Monomer predicts correctly but AfCycDesign predicts wrongly. (A) 2mw0. (B) 2m77.

#### 2.1.2 Performance comparison in disulfide bridge structure prediction

Disulfide bridges play an important role in cyclic peptide structures. However, AfCycDesign only focuses on the head-to-tail structure and neglects the existence of this fold type. As a result, there are instances where disulfide bridges are not connected or incorrectly connected within some structures in AfCycDesign. This lack of constraint introduces a vulnerability in structural accuracy. To avoid such case, HighFold_Monomer modifies the sequence separation between each pair of bonded cysteines to be one, which constrains disulfide connectivity, therefore correct disulfide bridge connectivities are formed for classes of structure with disulfide. In the case where exist multiple cysteines, our model enumerates all possible disulfide bridge pairs assembles in the input matrix and predicts corresponding folds. Constraints are placed on disulfide connectivity, rather than entrusting the network with the determination of these connections, ensuring that HighFold could avoid the situation of incorrect pairing of disulfide bonds. Then, our model chooses the top-5 predicted model based on pLDDT scores. This sufficient enumeration helps our HighFold_Monomer finds the best model.

As is shown in Figure 2C, HighFold_Monomer has a better ability to deal with various cyclic peptide structures with different disulfide bridges compared to AfCycDesign. Especially in predicting structures containing multiple disulfide bridges, HighFold_Monomer always gives biologically reasonable folding structures with correct disulfide bridge pairs. For example, the HighFold_Monomer can achieve the average RMSD value of 1.720 Å and AfCycDesign gets 3.256 Å in predicting cyclic peptide conformation including two disulfide bridge connections. The matrixes of different disulfide bond pairs are also put in Figure 2C. Figure 4 displays some examples that HighFold_Monomer predicts correctly but AfCycDesign predicts wrongly. In the predictions of 7l53 and 7m3u, AfCycDesign both generate inaccurate conformations with the RMSD values of 4.327 Å and 4.827 Å respectively. As we mentioned above, the disulfide bridges between cysteine residues play an important role in peptide structures. The disulfide bridge pair errors may lead to the total structure being wrong. In contrast to the AfCycDesign, our model can find the correct disulfide pairs and produce cyclic peptide structures with high accuracy. The RMSD values to 7l53 and 7m3u native conformations can achieve 2.305 Å and 1.058 Å, which is far superior to that of AfCycDesign.

**Figure 4.**
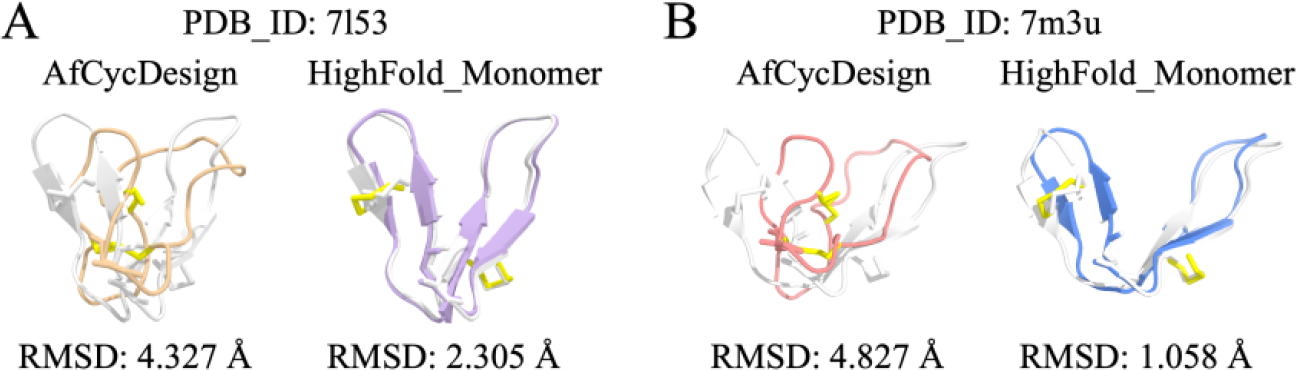
Disulfide bridge structures that HighFold_Monomer predicts correctly but AfCycDesign predicts wrongly. (A) 7l53. (B) 7m3u.

#### 2.1.3 Performance comparison in different secondary structures prediction

The network displays disparities in the prediction of structures with different secondary structures. Compared with the AfCycDesign, the predictive performance of High-Fold improves for each secondary structure category, especially for structures with β-sheet presence. Within structures characterized by α-helical and β-sheet, constraints on disulfide connectivity can affect the spatial positions of more amino acids. While in structures comprised solely of the random coil, disulfide connectivity constraints can only affect amino acids proximal to the corresponding cysteine, with limited impact on the spatial positions of more amino acids. Therefore, the network performs better in predicting the presence of α-helical and β-sheet structures, reaching an average of 1.287 Å and 1.478 Å respectively (Figure 2D). Figure 5 provides some examples to observe visually the ability gap between those models in secondary structure prediction. Take the 1df6 as an example, the HighFold_Monomer can give a more accurate random coil fold (RMSD 1.039 Å) with the constrain of the disulfide bond matrix, even though this secondary structure is very flexible.

**Figure 5.**
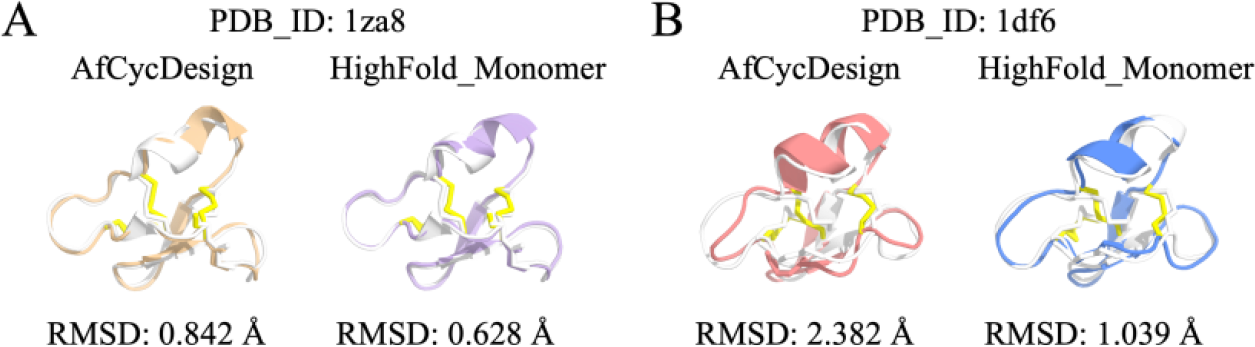
Secondary structures that HighFold_Monomer predicts correctly but AfCycDesign predicts wrongly. (A) 1za8. (B) 1df6.

#### 2.1.4 Performance comparison in structures prediction with different input lengths

We are interested in the performance of HighFold_Monomer when faced with different input lengths. As demonstrated in Figure 2E, HighFold has a better result in structures with shorter sequence lengths, and the most significant improvement in prediction accuracy was observed in structures with sequence lengths less than 30. The network performs best in predicting structures with sequence lengths of less than 20, with an average RMSD of 1.389 Å. For structures with sequence lengths greater than 30, the proportion of random coil regions in the structure increases, and the proportion of α-helical and β-sheet decreases, which likely explains a decrease in prediction accuracy.

#### 2.1.5 Correlation analysis between pLDDT and RMSD

PLDDT is the main metric to rank the predicted models given by HighFold_Monomer. We wonder if there exists a correlation between pLDDT and RMSD. To solve this puzzle, we make the correlation analysis between them, and the results are displayed in Figure 2F. There is no direct linear correlation between pLDDT and RMSD. However, we find that the higher pLDDT, the smaller the RMSD. Namely, the predicted cyclic peptide conformation is more likely to be close to the native structure with a higher pLDDT score. This finding may help readers to better assess the predicted results provided by HighFold_Monomer.

### 2.2 The performance of HighFold_Multimer in cyclic peptide complex structure prediction

In this section, we adapt a benchmarking data (detailed information can be found in the Method) to evaluate the performance of HighFold_Multimer in cyclic peptide complex structure prediction. Additionally, we compare our model with ADCP^10^, a famous CADD tool for docking cyclic peptides to proteins. In alignment with the methodology pursued by ADCP, we apply the F_nat_ as the principal metric to gauge the efficacy of models.

#### 2.2.1 Fnat comparison between HighFold_Multimer and ADCP

Figure 6A displays the performances of HighFold_Multimer and ADCP in cyclic peptide complex structure prediction. The computed average F_nat_ value for High-Fold_Multimer is 0.813, while that for ADCP stands at 0.415. As exemplified by the case of 1sfi, this HighFold_Multimer achieves a F_nat_ score of 0.930 when faced with such complex input where target and ligand lengths are 245, 14 respectively. In contrast, ADCP only gets an inferior F_nat_ value of 0.19, underscoring the superior performance of HighFold_Multimer in this scenario. The predicted performance difference is also very evident in the prediction of 4kel. The F_nat_ score of HighFold_Multimer is 0.846 and that of ADCP only achieves 0.24. Some structures remain challenges for both High-Fold_Multimer and ADCP. For example, HighFold_Multimer gets a low F_nat_ score of 0.584 when predicting the fold of 3p8f, which doesn’t mean a satisfying result. The ADCP performs worse than our model and only reaches an extremely low F_nat_ score of 0.04. To demonstrate the difference between those two models, we put a F_nat_ distribution in Figure 6C. As illustrated in this Figure, the F_nat_ distribution of HighFold_Multimer is higher than that of ADCP, further highlighting the superiority of our model in predicting cyclic peptide complexes.

**Figure 6.**
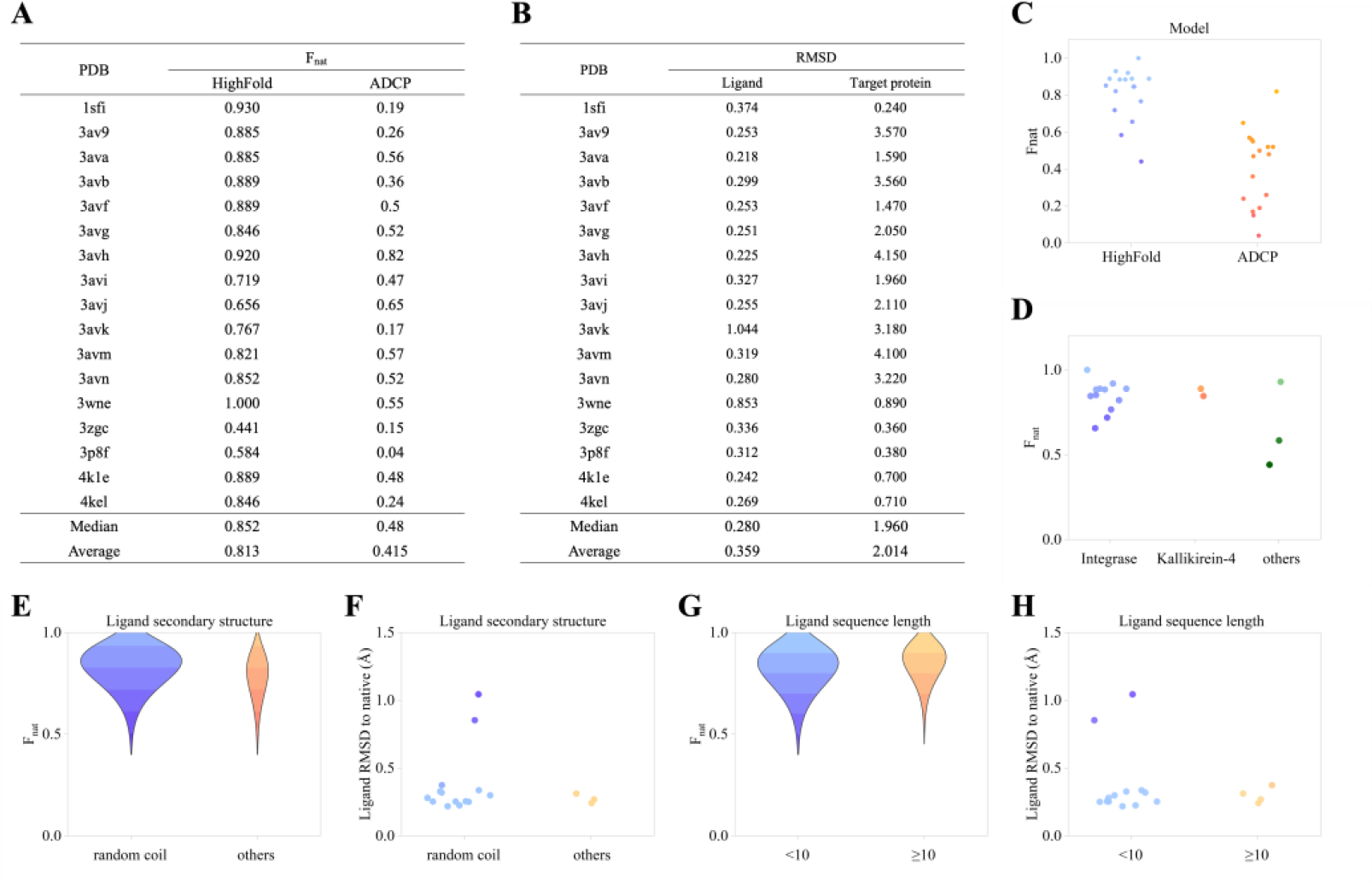
The results of models in cyclic peptide complex structure prediction. (A) F_nat_ results of HighFold_Multimer and ADCP. (B) RMSD results of ligands and target proteins within cyclic peptide complex structures generated by HighFold_Multimer. (C) F_nat_ distribution of HighFold_Multimer and ADCP. (D) F_nat_ distribution of HighFold_Multimer based on different target proteins. (E) F_nat_ distribution of HighFold_Multimer based on different secondary structure motifs of ligands. (F) RMSD distribution of ligands generated by HighFold_Multimer based on different secondary structure motifs. (G) F_nat_ distribution of HighFold_Multimer based on different input lengths of ligands. (H) RMSD distribution of ligands generated by HighFold_Multimer based on different input lengths of ligands. Note that all data analyses are based on top-1 results and the results of ADCP come from this paper.

What’s more, a noteworthy observation emerged about ADCP in this experiment. This approach exhibits a propensity towards generating random coil conformations of ligands, with little consideration of secondary structure motifs. However, HighFold-_Multimer usually give folds which are consistent with their native secondary structural features. In conclusion, HighFold_Multimer demonstrates a higher accuracy for predicting secondary cyclic peptide structures in comparison to ADCP.

#### 2.2.2 Performance in target protein structure prediction

As we mentioned above, the complex input contains protein and cyclic peptides. In this section, we focus on the HighFold_Multimer’s performance in target protein structure prediction. Those protein structures can be found in Figure 7. Figure 6B shows predicted target protein RMSD values to native protein structures. Even though the 3ava, 3avb, and 3avg contain the same protein target, their target protein RMSD is not relatively consistent, which are 1.590 Å,3.560 Å,2.050 Å. We guess it may be a result caused by multiple factors such stability of HighFold_Multimer, ligand size, and target protein’s structure characteristics.

**Figure 7.**
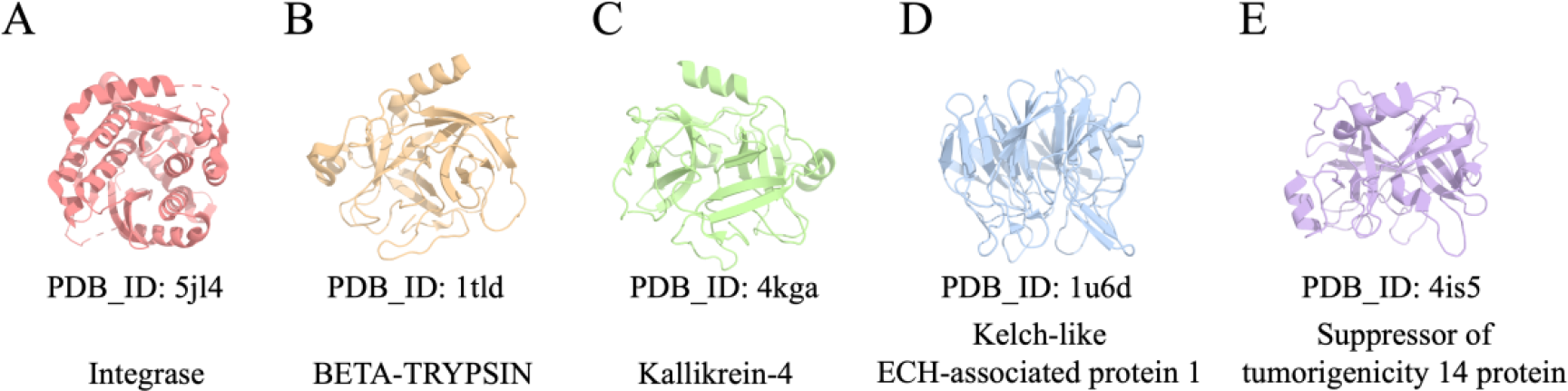
Target protein structures. (A) Integrase. (B) BETA-TRYPSIN. (C) Kallikrein-4. (D) ECH-associated protein 1. (E) Suppressor of tumorigenicity 14 protein.

In this work, we analyze the F_nat_ difference between different target proteins. As illustrated in Figure 6D, the HighFold_Multimer performs well in most predictions including target proteins. What’s more, we find integrase protein accounts for a large proportion of this dataset. Namely, the dataset of ADCP doesn’t have good diversity in a sense.

#### 2.2.3 Performance in ligand structure prediction

In this section, we discuss the performance of HighFold_Multimer in ligand structure prediction. Figure 6B illustrates the ligand RMSD difference of our model. As shown in Figure 6B, the RMSD of HighFold_Multimer’s predicted ligand can reach 0.218 Å at best, meaning this ligand model is close to the native conformation. Notably, the RMSD values of ligands span a range from 0.218 Å to 1.044 Å, and the overall average attains an impressive 0.359 Å, demonstrating that our HighFold_Multimer is good at dealing with ligand structure prediction in most cases.

In addition, we discuss the performance of HighFold_Multimer based on secondary structure motifs of ligands. However, there are no evident differences between model performances based on the random coil and other secondary structure motifs. Figure 6E and F shows the performance difference of the model in different secondary motif ligand predictions. There are not many structures of ligands containing secondary motifs in this dataset and it seems that HighFold_Multimer can deal with most cyclic peptide structure prediction in our experiment. Figure 8 shows some structure comparison between that of HighFold_Multimer and that of the traditional experiment. Although the loop structure of 3wne is flexible, our model can get the F_nat_ value of 1.000 and the ligand RMSD value of 0.853 Å. As for the 4kle where β-sheet exists, our model can achieve the F_nat_ value of 0.889 and the RMSD value of 0.242 Å. Anyway, our model performs well in this benchmarking dataset.

**Figure 8.**
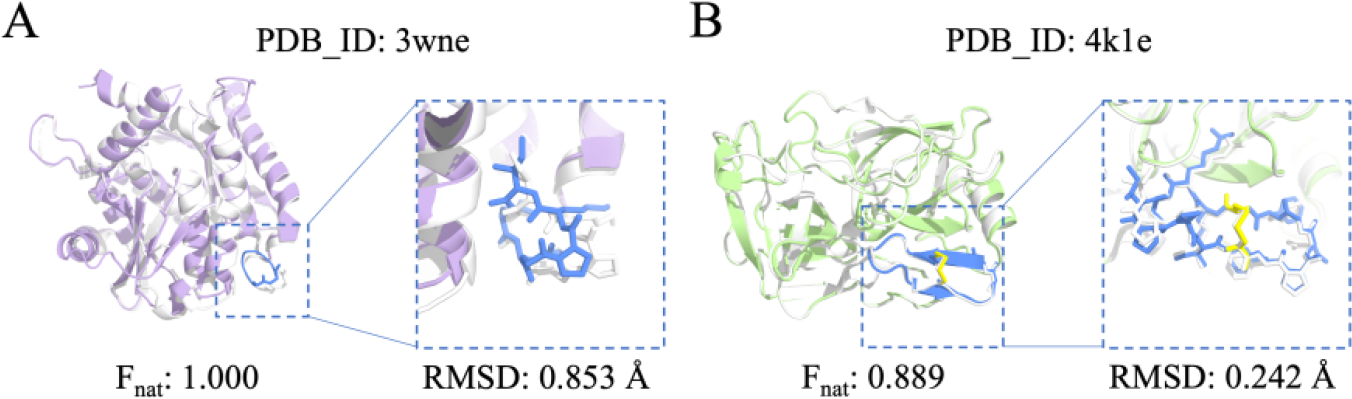
Structure examples with different ligand secondary structure motifs. (A) 3wne where ligand only includes random coil structure. (B) 4kle where ligand includes random coil and β-sheet structures.

#### 2.2.4 Performance in structures prediction with different ligand’s input lengths

We also explore the influence of ligand input length in this task and the results can be found in Figures 6G and 6H. Due to the limitation of the dataset, there are not many structures with ligand input lengths that compass 10. No matter whether in F_nat_ or RMSD distributions, there is no clear correlation between predicted structures with ligand input lengths. As shown in Figure 9, the ligand RMSD can both reach around 0.3 Å even though their sizes are different.

**Figure 9.**
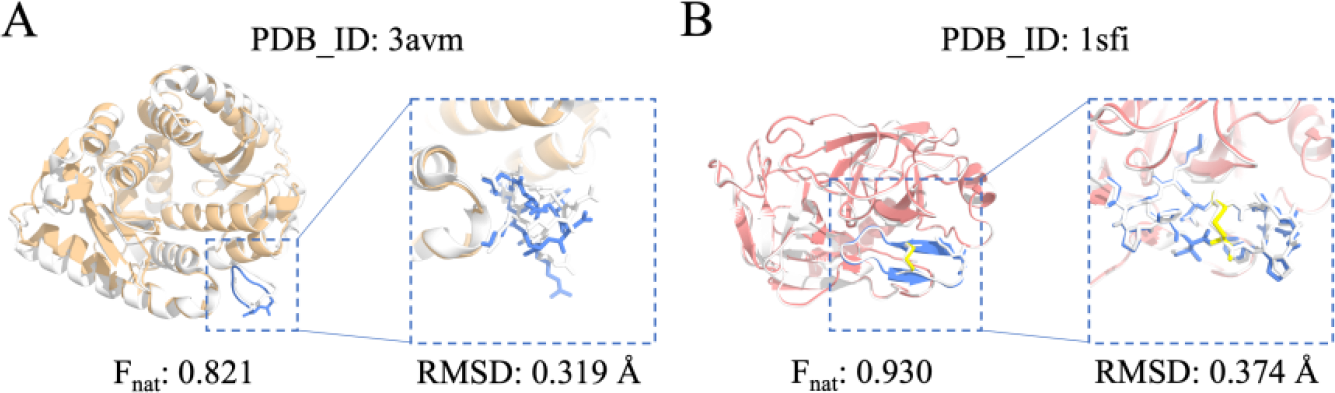
Examples of cyclic peptide complex structures with different ligand input lengths. (A) predicted structure of 3avm with ligand input length of 8. (B) predicted structure of 1sfi with ligand input length of 14.

### 2.3 The performance of HighFold_Multimer in external cyclic peptide complex structure prediction

To further demonstrate the generality of our DL-based HighFold_Multimer, we make a cyclic peptide complex prediction experiment based on an external dataset. Detailed information on this data can be found in the Method. Figure 10 illustrates the power of HighFold_Multimer in predicting various cyclic peptide complex folds. This model performs best in predictions of 6u22 and 6bvh, and their F_nat_ are both 0.948, almost one. Compared to the cyclic peptide complex data of ADCP, our external dataset is more diverse. The lengths of cyclic peptide ligands range from 6 to 34 and there are different secondary structures in native conformations. It should be noted that ADCP only deals with cyclic peptide ligands with lengths up to 20. However, there is no length limitation in our model. Figure 10 also displays the predicted models and native structures that cyclic peptide ligand lengths are 14, 34, 14. The RMSD values of those ligand structures are 0.359 Å,0.230 Å and 0.431 Å respectively, meaning a fact that High-Fold_Multimer can deal with such macrocyclic peptide complexes well. All results of our work can be found in supporting information.

**Figure 10.**
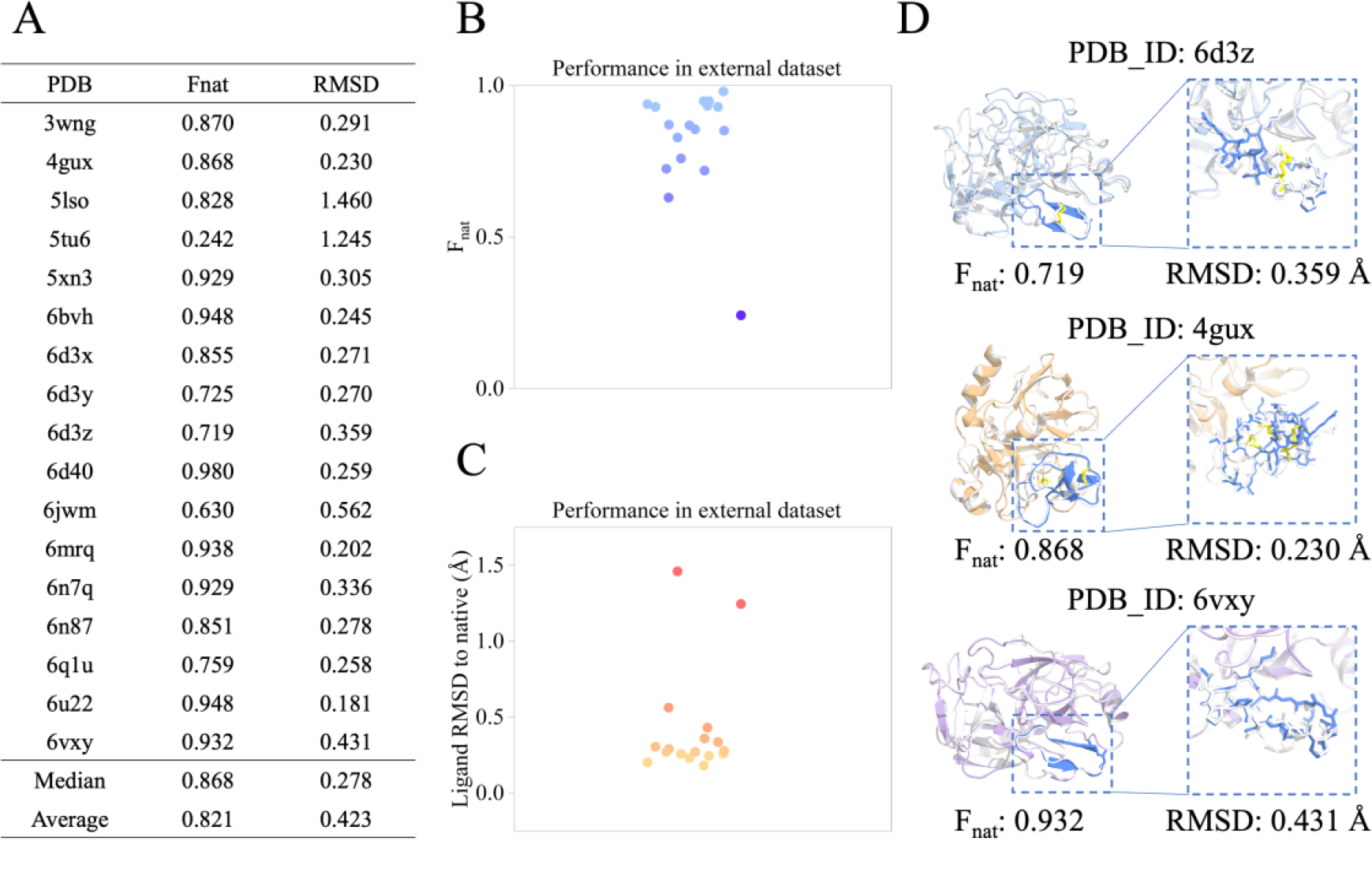
The results of HighFold_Multimer in external cyclic peptide complex structure prediction. (A) F_nat_ and ligand RMSD results of HighFold_Multimer. (B) F_nat_ distribution of High-Fold_Multimer (C) ligand RMSD distribution of HighFold_Multimer. (D) Representative predicted complex structures generated by HighFold_Multimer.

## 3. CONCLUSION

In this work, we propose the HighFold framework, where two important terms are included in the structure prediction of cyclic peptides and their complexes respectively. The trade-off between accuracy and efficiency enables our models to become a potential tool for further fitting the studies in related areas. Specifically, our models can both retain the outstanding backbone and disulfide bridges sidechain folds of cyclic peptides in monomers and complexes, which are comparable or even superior to other baselines. We further emphatically explore the effects of several important features of cyclic peptide structures on HighFold’s performance. The results indicate that this model is better at dealing with cyclic peptides involving different lengths and secondary structures. We believe our work could serve as a reliable tool for cyclic peptide-related structure prediction, and our deep analysis could also provide valuable insights into the development of these peptide classes.

## 4. METHOD

### 4.1 Datasets

#### 4.1.1 Cyclic peptide monomers set

The cyclic peptide monomers set comes from the work of Bhardwaj et al. There are 63 structures with lengths from 12 to 39. In this data, all structures are experimentally investigated and their secondary structure includes 7 α-helical and 37 β-sheet,10 random coil, and 9 mixed structures.

#### 4.1.2 Cyclic peptide complexes set

The cyclic peptide monomers set comes from the work of ADCP. There are 17 structures with ligand lengths from 6 to 14 in this data and the secondary structure motifs of ligands cover 3 β-sheet and 14 random coil structures.

#### 4.1.3 External cyclic peptide complexes set

Cyclic peptide complexes in an external dataset are downloaded from the PDB database. This data contains 17 structures with ligand lengths from 6 to 34, and the secondary structure motifs of ligands cover 10 β-sheet, 5 random coil and 2 mixed structures.

### 4.2 Model

In this section, we introduce the details of our proposed HighFold and HighFold-Multimer, including the construction of the offset matrices, generation of the disulfide bridge’s combinations, and additional processing for HighFold-Multimer r, as shown in Figure 11.

**Figure 11.**
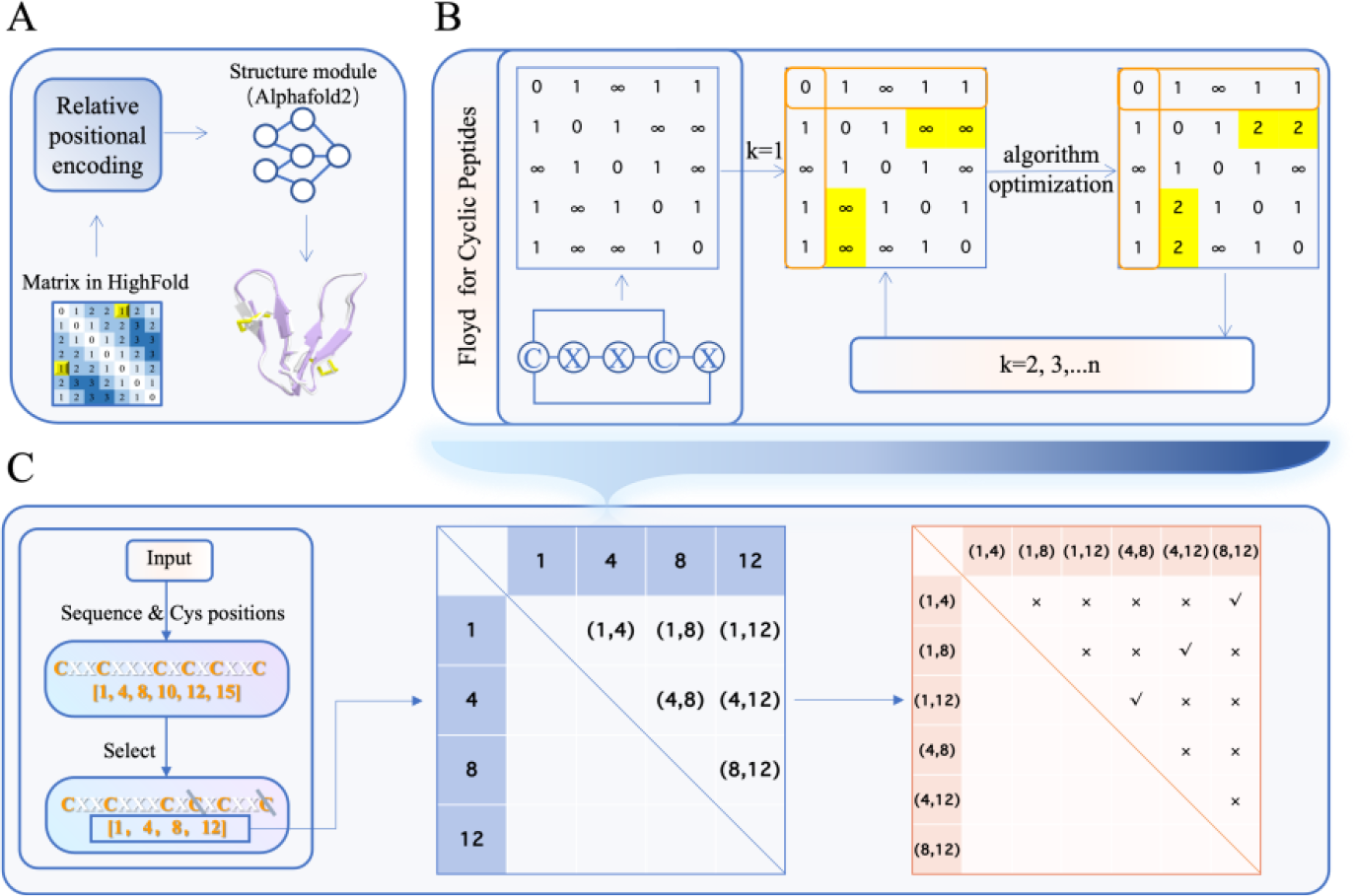
The algorithms in HighFold. (A) The basic flow of HighFold. (B) Floyd for Cyclic Peptides algorithm for CPCM matrix construction. (C) The calculation flowchart of disulfide bridges’ combinations. In subfigure (B) and subfigure (C), “C” denotes Cysteine, and “X” denotes any amino acids except Cysteine.

#### 4.2.1 Cyclic Peptide Connectivity Matrix

The original AlphaFold processes the distance information between amino acids in the linear mode. However, the performance will drop dramatically when it comes to the situation of cyclic peptides due to inaccurate distance information, especially in the case of multiple cycles by different cyclization mechanisms. Here we propose a dynamic programming algorithm, namely Floyd for Cyclic Peptides (FCP) algorithm, to enhance the distance information of each pair of amino acids by additionally constructing a positional offset matrix for each cyclic peptide. The FCP algorithm is based on the classical Floyd’s algorithm^26^, which is proposed to find the shortest paths in a directed weighted graph in the approach of dynamic programming.

In particular, given the sequence of one cyclic peptide with N amino acids *a*_1_*a*_2_ … *a*_*N*_, the positional offset matrix can be constructed by calculating the shortest paths between each amino acid. Initially, the adjacency matrix is constructed according to the information of head-to-tail cyclization, disulfide bridge conformation, and the natural connectivity information between two adjacent amino acids. Then similar to the Floyd algorithm, the FCP algorithm conducts three nested loops, that is, the outer loop, the middle loop, and the inner loop. Each loop runs through the number of amino acids. The key point is taking the amino acid in the outer loop as an intermediate point recursively, and updating the distance of all the pairs of amino acids if the distance via the intermediate point is shorter than the original one. In summary, the procedure of the FCP algorithm is described as Algorithm 1. Note that the FCP algorithm is also suitable for the scenarios of linear peptides by giving the empty set of disulfide bridges and commenting out the head-to-tail cyclization.

[Offset matrix construction by Floyd’s algorithm]

##### Algorithm 1 Floyd for Cyclic Peptides (FCP) algorithm

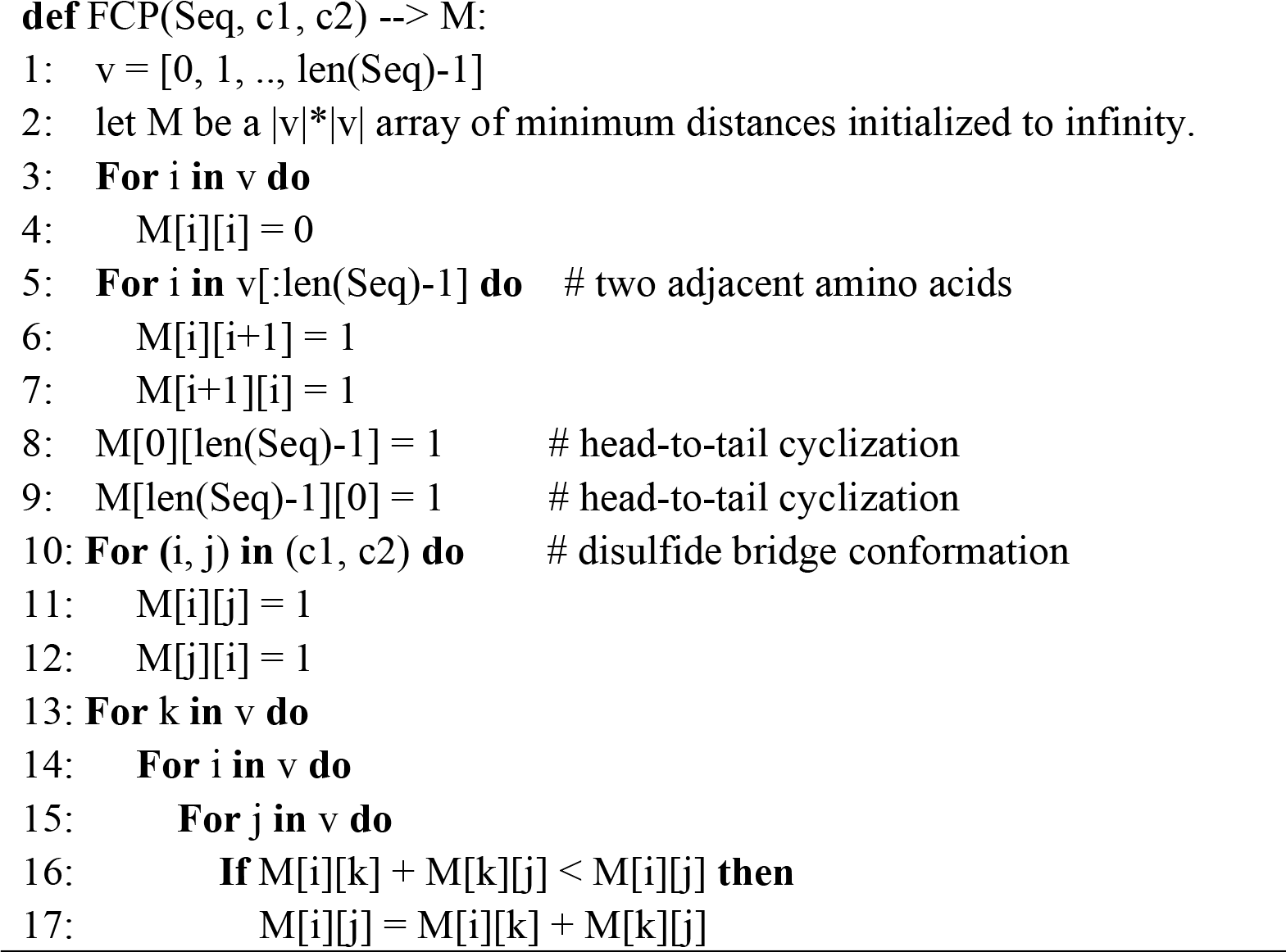

#### 4.2.2 Combinations of Disulfide Bridges

In practice, given a sequence of amino acids, multiple disulfide bridge combinations can be constructed if there are more than two cysteines in the sequence of amino acids. Take three cysteines a, b, and c as an example, there are three different pairs <a, b>, <a, c> and <b, c> to build the disulfide bridges. Here we propose a combinatorics-based algorithm, namely the Complete Set Of Disulfide Bridges (CSDB) algorithm, to enumerate all possible disulfide bridge combinations, and its procedure is described as Algorithm 2.

First of all, find the positions of all cysteines in the amino acid sequence, written as vector v=[c_1, c_2, …, c_N], where N denotes the number of cysteines. Then two loops are employed to generate the disulfide bridge’s combinations, namely the outer loop and the inner loop. The outer loop enumerates all even numbers that are less than or equal to n and feeds each even number to the inner loop. The reason why we choose the even numbers only is that all the disulfide bridge combinations for the case of odd numbers can be completely covered by the case of even numbers. The inner loop aims to generate the disulfide bridge’s combinations according to the given even number of cysteines. Specifically, they usually include several steps, that is, cysteine pair construction, cysteine pair combination generation, and cysteine pair combination validity verification. For example, given an even number k, let v_k=[1,2,…,k] denotes the index of cysteines, then cysteine pairs are constructed by conducting a 2-ary Cartesian power operation over vector v_k, which is formulated as C_pair = {(i, j) | I in v_k, and j in v_k}. Similarly, cysteine pair combinations are generated by conducting a k/2-ary Cartesian power operation over the set of cysteine pairs C_pair, which results in a set of cysteine pair combinations, S_comb = {(x_1,x_2,…,x_{k/2}) | x_1 in C_pair, x_2 in C_pair, …, and x_{k/2} in C_pair}. However, the combinations with overlap cysteines are invalid, because one cysteine cannot be in different disulfide bridges at the same time in a fixed cyclic peptide structure, so we simply remove these invalid combinations from S_comb subject to that neither of x_1[0], x_1[1], …, x_{k/2}[0] and x_{k/2}[1] are equal to each other.

[Generation of disulfide bridge’s combinations]

##### Algorithm 2 Complete Set of Disulfide Bridges (CSDB) algorithm

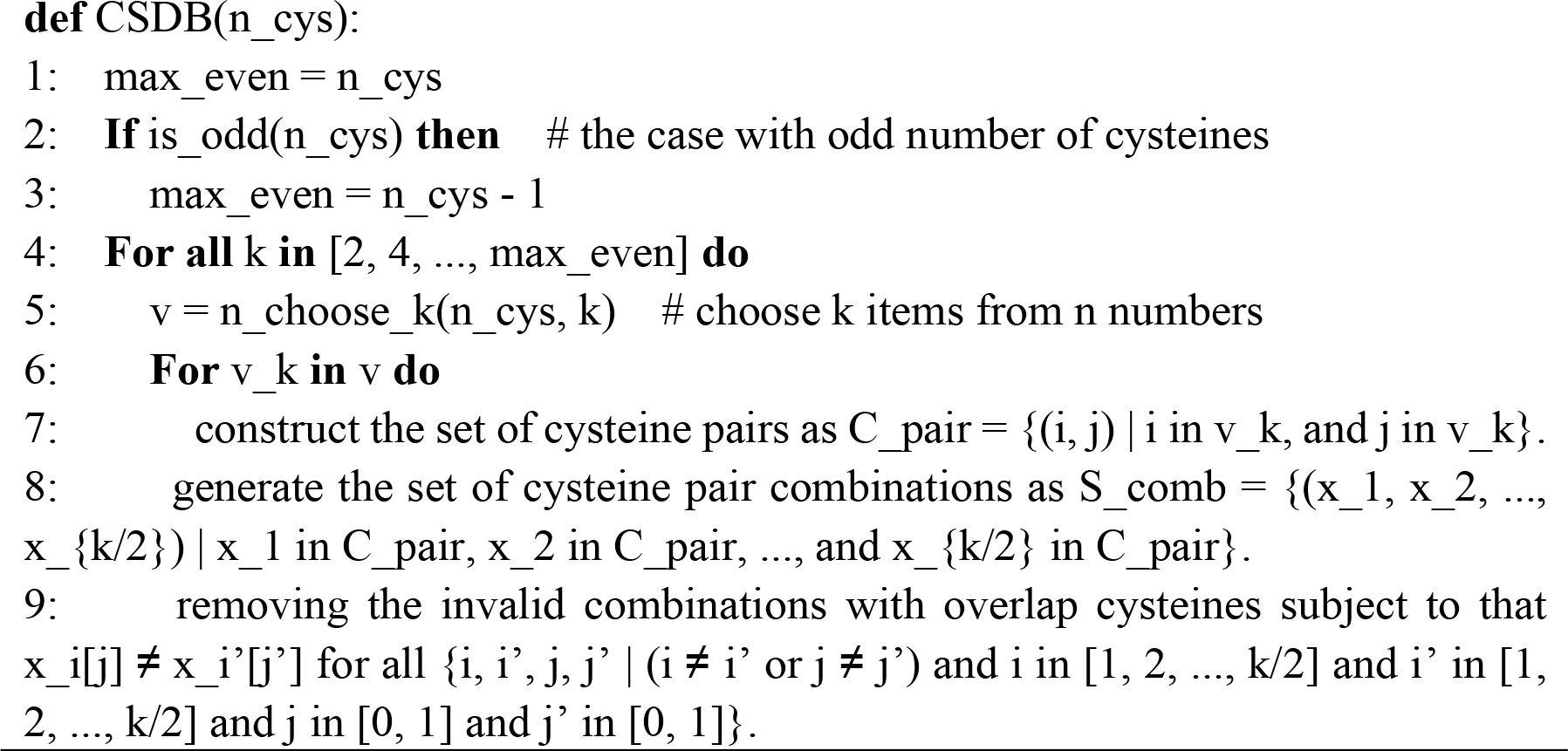

#### 4.2.3 Additional Processing with HighFold-Multimer

When it comes to the scenarios of cyclic peptide complex, the positional offset matrix can be built by explicitly assembling the positional offset matrices of each cyclic peptide monomer in HighFold. Supposing that the cyclic peptide complex contains N amino acid sequences with length L=[l_1, l_2, …, l_N] and the corresponding positional offset matrices S_M={M_i | i in [1, 2, …, N]} respectively. The positional offset matrix for cyclic peptide complex can be formulated as a |sum(L)|*|sum(L)| block diagonal matrix M = block-diagonal(M_1, M_2, …, M_N), where each diagonal block is a positional offset matrix of one cyclic peptide monomer, and it can be written in the matrix format as follows,

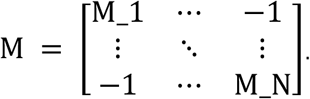

Note that elements ‘-1’ at non-diagonal blocks of M do not denote the distances, actually they are just some indicators to distinguish different amino acid chains/sequences.

### 4.3 Metrics

In this section, we evaluate the performance of HighFold from several different perspectives, including RMSD, F_nat_, pLDDT, and M_conf_, where RMSD and F_nat_ are used to evaluate model effectiveness, and pLDDT and M_conf_ are used as criteria for model selection.

RMSD, which is a classical measure of the average error between the predicted values and the observed values, here is employed to evaluate the average distance between the atom coordinates predicted by an estimator and the corresponding atom coordinates from chemical experiments. Specifically, the calculation of RMSD can be formulated as

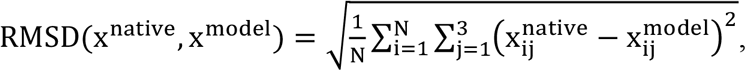

where x^native^ denotes cyclic peptide conformation from chemical experiments, x^model^ denotes cyclic peptide conformation predicted by the estimator, index i traverses the number of atoms N, and index j traverses the 3D coordinates. F_nat_^27^ is to measure the recall of native interfacial contacts preserved in the interface of the predicted cyclic peptide complex, and its calculation can be formulated as

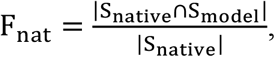

and

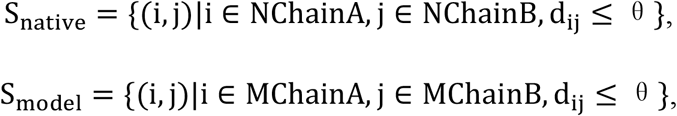

where Snative denotes the set of atom pairs with a distance less than threshold θ between the receptor and the ligand in the experimental structure, Smodel denotes the set of atom pairs with a distance less than threshold θ between the receptor and ligand in the predicted structure, operator ∩ denotes the intersection of two sets, operator |.| denotes the size of a set. Note that the distance threshold θ can be set to different values, and here we set it 5Å in our experiments. Generally, we employ different cutoffs on the F_nat_ measure to assign predicted conformations into the four quality classes: Incorrect (Fnat < 0.2), Acceptable (0.2 < F_nat_ < 0.5), Medium (0.5 < F_nat_ < 0.8), or High (0.8 < Fnat).

pLDDT^15^, an abbreviation for predicted local distance difference test, is a confidence of the predicted structure by evaluating the difference of local distance. Take an amino acid as an example, its calculation can be formulated as

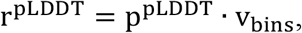

where P^PLDDT^ = softmax(f(s)) denotes the output of cascade operations, that is, mapping the amino acid’s embedding to a |v_bins_| dimensional space and the softmax of the intermediate results, and v_bins_ is the bin vector.

Mconf is used as a model confidence metric in the evaluation of HighFold-Multimer, which is a weighted combination of pTM and ipTM to take into consideration TM-score^28^ of both the entire sequence and the interaction interface. Its calculation can be formulated as

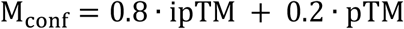

where pTM denotes predicted TM-score of the entire sequence, which is a confidence of TM-score based on the predicted structure. Similarly, ipTM denotes predicted TM-score of the interaction interface only.

## Supporting information

results

## 5. ACKNOWLEDGMENT

This project was supported by the Natural Science Foundation of Zhejiang Province (LD22H300004).

